# Deconvolved tumor adipocyte proportions and high grade serous ovarian carcinoma survival

**DOI:** 10.64898/2026.01.14.699527

**Authors:** Adriana Ivich, Laurie Grieshober, Natalie R. Davidson, Grace Y. Akatsu, Lauren C. Peres, Stephanie C. Hicks, Jeffrey R. Marks, Joellen M. Schildkraut, Jennifer A. Doherty, Casey S. Greene

## Abstract

**Background:** Single-cell-based analyses of high-grade serous ovarian carcinoma (HGSOC) survival have largely ignored adipocytes, which are fragile and under-represented in single-cell references. Adipocytes are known active components of the tumor microenvironment in many cancers, and HGSOC tumors frequently metastasize to the omentum, a lining of adipose tissue.

**Methods:** We created a composite reference that combines single-nucleus adipose profiles with published HGSOC single-cell data to deconvolve 588 bulk RNA-seq tumours from the *Schildkraut* cohorts. We used stage-stratified Cox models to quantify the association between intratumoural adipocyte fractions and overall survival while adjusting for age, body mass index (BMI), race, and residual disease. We also evaluated associations with deconvolved immune, stromal, and epithelial cell groups.

**Results:** A 10% increase in estimated tumor adipocyte content was associated with a 41% increase in the hazard of death (HR = 1.41, 95% CI 1.18-1.70, p = 0.0002) after adjusting for age, BMI and race (n=566). A 10% increase in immune cell proportion was associated with favorable survival (HR = 0.82, 95% CI 0.69-0.97, p = 0.024). Stromal and epithelial macro-fractions were not associated with survival. Associations with adipocyte and immune cell type proportions were unchanged in models additionally controlling the other cell type proportions. Results were similar after additionally adjusting for residual disease after debulking surgery.

**Conclusions:** Adipocytes may be a tumor-intrinsic factor associated with adverse outcomes in HGSOC. Quantifying adipocyte burden using bulk RNA-seq could enhance risk stratification and guide the development of adipocyte-targeted therapies.

## Introduction

High-grade serous ovarian cancer (HGSOC) accounts for 70-80% of all epithelial ovarian cancers (EOC) [1, 2]. Only about half of patients survive five years [2]. The majority of HGSOC are derived from the epithelium of the fallopian tube and ovary [1], and most patients have metastasis in the omentum, a lining of adipose tissue in the peritoneal cavity, by the time of diagnosis [1].

Tothill et al., and later The Cancer Genome Atlas (TCGA), identified four HGSOC molecular subtypes: immunoreactive, differentiated, proliferative, and mesenchymal, each with distinct survival outcomes [3, 4]. While subsequent studies have produced varying results regarding the number and definition of HGSOC subtypes [5–7], these studies have consistently reported associations between subtype classification and survival outcomes, suggesting a true underlying biological mechanism [4, 6–10]. The immunoreactive subtype has the best prognosis, generally followed by differentiated and proliferative with intermediate outcomes, while the mesenchymal subtype consistently shows the poorest survival. Studies examining molecular features of HGSOC often lack representation across racial groups, even though survival disparities exist, with Black individuals experiencing poorer outcomes than White individuals [11, 12]. Challenges in interpreting subtypes from bulk profiling have shifted attention toward the tumor micro-environment (TME) as a possible source of observed survival differences [7]. The TME is the community of non-cancerous cells that surrounds and shapes each HGSOC tumor, and the cell type composition can be either studied through a higher resolution transcriptomic assay, single-cell RNA-seq (scRNA-seq), or through bulk RNA-seq using deconvolution to estimate cell type proportions [13]. Immune cells, fibroblasts, endothelial cells, and other TME elements can promote or decrease tumor growth and affect drug response, and in turn affect patient prognosis [14]. In HGSOC, Hippen et al. reported that tumors rich in fibroblasts were associated with shorter survival, whereas immune-associated tumors fared better [15]. Yet most TME studies stop at immune or fibroblast signals and overlook adipocytes: the fat cells that dominate the omentum and may carry their own prognostic weight.

Growing evidence shows that adipocytes are key components of the TME in EOC, driving disease progression through multiple mechanisms [16–19]. Adipocytes actively interact with EOC cells and can profoundly influence tumor progression through immunosuppression, angiogenesis, proliferation, metastasis, and treatment resistance [16, 18, 20–22]. Despite the evidence of adipocytes’ influence in cancer progression, it is unclear whether adipocyte abundance in HGSOC tumors correlates with patient outcomes.

Studying adipocyte composition in HGSOC tumors presents its own challenge, since these large and delicate cell types are often lost in scRNA-seq data and thus are not included in deconvolution predictions [23]. We previously evaluated strategies to incorporate single nucleus RNA-seq (snRNA-seq) references alongside scRNA-seq references, allowing us to estimate adipocyte proportions in HGSOC samples from bulk RNA-seq samples [24].

In this study, we examine whether adipocyte proportions in HGSOC tumors are related to poorer survival outcomes using deconvolution to estimate cell proportions in a previously characterized bulk RNA-seq dataset from the *Schildkraut* cohorts [12]. To estimate adipocyte proportions, we use a snRNA-seq dataset of adipose tissue along with scRNA-seq HGSOC datasets to have a complete reference with all TME components. We use Cox Proportional Hazards (CPH) models to estimate associations between adipocyte proportions and survival outcomes, taking into account covariates such as stage, age, body mass index (BMI), and presence of residual disease after debulking surgery.

We also explore whether the proportions of other deconvolved major cell groups - immune cells, stromal cells (including fibroblasts and endothelial cells), and epithelial cells (including tumor cells) - were associated with survival outcomes. Additionally, we examined whether clinical covariates were associated with differences in cell-type proportions, and whether cell-type proportions were associated with transcriptomic subtypes. This exploratory analysis aimed to uncover broader TME-related patterns in prognosis that may inform future mechanistic or therapeutic studies.

## Results

### Tumor adipocytes and survival

We leveraged bulk RNA-seq data from the 588 HGSOC tumor samples from the *Schildkraut* cohorts [12] (Table 1). To estimate the cell composition of the tumor samples, we used bulk RNA-seq deconvolution on the raw count profiles using a cell reference that combined published HGSOC scRNA-seq datasets [25] with adipose tissue snRNA-seq datasets [23] (Fig. 1a). For the *Schildkraut* samples, a pathologist reviewed slides to select areas with tumor tissue for downstream analyses, so the RNA-seq data primarily represented cell types that are within the tumor, and not the surrounding tissue. The obtained estimated cell-type proportions for all samples are shown in Supplemental Fig. 1.

**Fig. 1.**
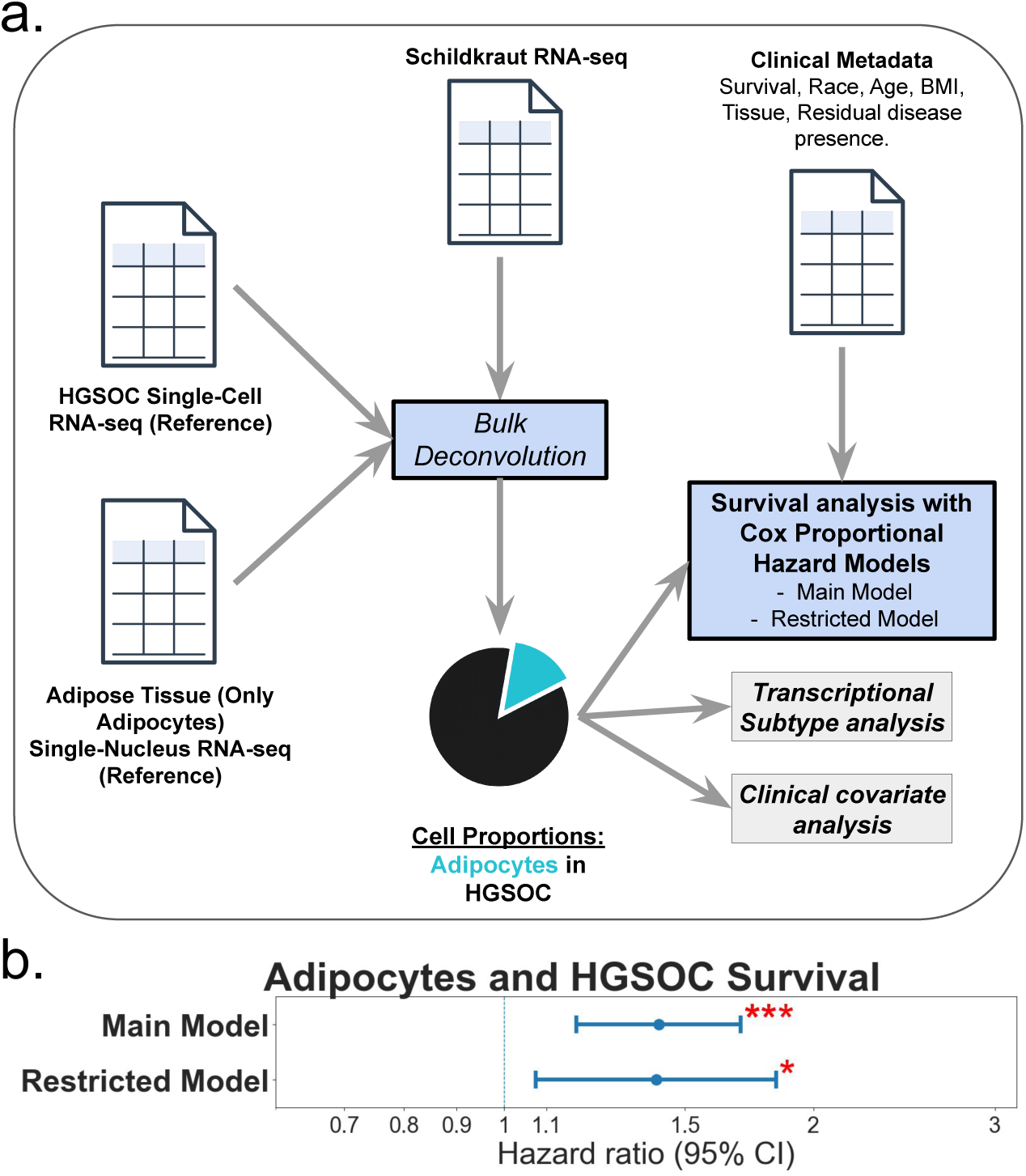
Adipocyte proportion quantification and association with high-grade serous ovarian cancer (HGSOC) Survival. The figure illustrates the methodology for quantifying adipocyte proportions in HGSOC and the subsequent survival analysis. Panel a shows the workflow: HGSOC scRNA-seq data and snRNA-seq data from adipose tissue (only adipocytes) serve as reference datasets. These, along with the *Schildkraut* bulk RNA-seq data, are used for bulk deconvolution to predict cell proportions (tumor composition), represented by a pie chart where a turquoise slice indicates the adipocyte proportion. Both the calculated adipocyte proportions and the clinical data are then used in the survival analysis using CPH modelling. Panel b presents a forest plot of the HRs for a 10% increase in adipocyte proportions and HGSOC survival for the main (stage-stratified and adjusted for age, BMI, and race, n = 566: HR = 1.41, 95% CI 1.18-1.70) and adjuvant therapy-restricted models (additionally adjusted for residual disease, n =262: HR = 1.41, 95% CI 1.07-1.84). Red asterisks indicate statistically significant associations (* p < 0.05; ** p < 0.005; *** p < 0.0005). The dashed vertical line at HR = 1 serves as a reference for no association. BMI; body mass index. scRNA-seq; single-cell RNA sequencing. snRNA-seq; single-nucleus RNA sequencing. HGSOC; High-Grade Serous Ovarian Carcinoma. CPH; Cox Proportional hazards. HR; Hazard Ratios.

**Table 1.** Characteristics of the *Schildkraut* HGSOC cohort The main models include individuals with complete information on BMI and FIGO stage (n=566), and the restricted models include individuals who received adjuvant chemotherapy and who had information on debulking status (n=262). BMI: body-mass index. FIGO: International Federation of Gynecology and Obstetrics.

| Clinical Variable | Category | Number of Patients |  |  |
| --- | --- | --- | --- | --- |
|  |  | All (n=588) | Main Model (n=566) | Restricted Model (n=262) |
| Race | Black | 272 (46.3%) | 263 (46.5%) | 151 (57.6%) |
|  | White | 316 (53.7%) | 303 (53.5%) | 111 (42.4%) |
| Age at Diagnosis | Age | n=588, mean=58.22, SD=9.65, min=26, max=78 | n=566, mean = 58.25, SD=9.53, min=1, max=78 | n=262, mean=58.32, SD=9.59, min=36, max=78 |
| Vital Status | Alive/Censored | 117 (19.9%) | 110 (19.4%) | 61 (23.3%) |
|  | Deceased | 471 (80.1%) | 456 (80.6%) | 201 (76.7%) |
| Years from interview to last follow up | Number of years | mean=5.73, SD: 4.89, min=0.11, max=20.96 | mean=5.73, SD=4.88, min=0.11, max=20.96 | mean=6.17, SD=4.84, min=0.11, max=20.96 |
| FIGO Stage | I | 48 (8.2%) | 47 (8.3%) | 17 (6.5%) |
|  | II | 39 (6.6%) | 38 (6.7%) | 23 (8.8%) |
|  | III | 469 (79.8%) | 459 (81.1%) | 211 (80.5%) |
|  | IV | 23 (3.9%) | 22 (3.9%) | 11 (4.2%) |
|  | Unknown | 9 (1.5%) | 0 | 0 |
| BMI (1 year prior to diagnosis) | BMI (kg/m2) | mean=29.58, SD=7.78, min=14.81, max=74.39 | mean=29.33, SD=7.79, min=14.81, max=74.39 | mean=29.98, SD=8.19, min=14.81, max=74.39 |
|  | Unknown | 13 (2.2%) | 0 | 0 |
| Residual disease | No visible disease (0 mm) | 110 (18.7%) | 107 (18.9%) | 97 (37.0%) |
|  | Residual disease, diameter of largest residual tumor < 1 cm | 99 (16.8%) | 96 (17.0%) | 90 (34.4%) |
|  | Residual disease, diameter of largest residual tumor ≥ 1 cm | 91 (15.5%) | 90 (15.9%) | 75 (28.6%) |
|  | Unknown | 288 (48.9%) | 273 (48.2%) | 0 |
| Received adjuvant chemotherapy | Yes | 290 (49.3%) | 281 (49.7%) | 262 (100%) |
|  | No | 7 (1.2%) | 7 (1.2%) | 0 |
|  | Unknown | 291 (49.5%) | 278 (49.1%) | 0 |
| Anatomic location of tissue sample | Fallopian Tube and Ovary | 120 (20.4%) | 115 (20.3%) | 72 (27.5%) |
|  | Omentum | 16 (2.7%) | 16 (2.8%) | 12 (4.6%) |
|  | Other | 11 (1.9%) | 11 (1.9%) | 8 (3.0%) |
|  | Unknown | 441 (75.0%) | 424 (74.9%) | 170 (64.9%) |

We limited survival analyses to individuals with known BMI and International Federation of Gynecology and Obstetrics (FIGO) stage, resulting in 566 patients in total. After deconvolution, we multiplied the predicted adipocyte proportions (0-1) by 10 so that hazard ratios (HRs) reflect a 10% increase in adipocyte content, and we stratified all CPH models by FIGO stage to satisfy proportional-hazards assumptions. Results are reported for both a main model (stage-stratified, adjusted for age, BMI, and race, n = 566), and a restricted model (including only individuals who were known to have received adjuvant chemotherapy and additionally adjusted for residual disease, n = 262).

Adipocyte abundance was associated with increased mortality (Fig. 1b). In the main model, every 10-percentage-point rise in adipocyte content increased the hazard of death by 41% (HR = 1.41, 95% CI 1.18–1.70, p = 0.0002). The association was similar in the restricted model (HR = 1.41, 95% CI 1.07-1.84, p = 0.013). Results stratified by race were similar (Supplemental Fig. 2). A 10% increase in adipocyte abundance was associated with a 42% higher hazard of death in Black patients (HR = 1.42, 95% CI 1.08-1.87, p = 0.011, n = 263) and a 43% higher hazard in White patients (HR = 1.43, 95% CI 1.11-1.84, p = 0.005, n = 303) after stratifying by stage and controlling for age and BMI. Among individuals who received adjuvant chemotherapy, additionally controlling for residual disease showed comparable associations that did not reach statistical significance (Black: HR = 1.40, 95% CI 0.96-2.04, p = 0.079, n = 151; White: HR = 1.41, 95% CI 0.94-2.12, p = 0.099, n = 111).

Together, these findings suggest that intratumoural adipocytes capture a tumor-intrinsic feature that may independently be associated with worse survival in HGSOC, increasing mortality risk by about 40% for each 10% increment in adipocyte content within the tumors.

### Immune, stromal, and epithelial proportions’ relationship to survival

We performed an exploratory analysis to investigate whether the other TME cell groups identified using deconvolution were associated with survival. We mapped the cell types into three additional TME groups: immune, stromal, and epithelial (Supplemental Fig. 3a).

For each tumor, we summed the proportions for each cell type within each cell group and multiplied the result by ten, so a one-unit change in the CPH model represents a 10% rise in the compartment, mirroring the scaling used for adipocytes. We similarly fit two models for each cell group: a main model (stage-stratified and adjusted for age, BMI, and race), and a restricted model among individuals who received adjuvant chemotherapy, additionally adjusted for residual disease.

As expected, immune cell abundance was significantly associated with improved survival in the main model (HR = 0.82, 95% CI 0.69-0.97, p = 0.024), with a similar magnitude in the restricted model (HR = 0.80, 95% CI 0.61-1.03, p = 0.085) (Supplemental Figure 3b). Stromal and epithelial content were not markedly associated with survival (Supplemental Figure 3c-d).

To confirm that the adipocyte association was not confounded by other TME components, we performed a sensitivity analysis including adipocyte proportions together with immune and stromal fractions in the same models, excluding epithelial content to reduce collinearity (Supplemental Fig. 4). The results were consistent with the primary analysis: adipocyte abundance remained significantly associated with worse survival in both the main (HR = 1.42, 95% CI 1.17-1.71, p = 0.0003) and restricted (HR = 1.40, 95% CI 1.05-1.86, p = 0.021) models, while immune abundance retained a significant protective association in the main model (HR = 0.80, 95% CI 0.67-0.96, p = 0.014), with a similar magnitude in the restricted model (HR = 0.78, 95% CI 0.60-1.01, p = 0.059) (Supplemental Fig. 4). Stromal content again showed no association with survival. Both models exhibited borderline proportional-hazards violations, so these findings are presented for transparency rather than inference (Supplemental Fig. 4).

In summary, these analyses suggest that after controlling for the major cell types, increased adipocyte proportions are associated with increased mortality, and increased immune cell proportions are associated with more favorable survival.

### Cell-type composition by age and BMI

Further exploratory analysis aimed to evaluate whether the intra-tumour cell composition differed by age and BMI. We first correlated each macro-fraction or cell type group (adipocytes, immune, stromal, and epithelial) with continuous age and BMI using Spearman’s correlation (ρ), yielding eight tests in total. We then fit a β-regression to estimate the change in fraction produced by a one-standard-deviation rise in each cell type group.

None of the eight Spearman correlations (cell groups vs. age, cell groups vs. BMI) were statistically significant after Bonferroni adjustment (all *p-adj* ≥ 0.70) (Fig. 2a-b). The strongest raw signal was a weak inverse correlation between age and the immune fraction (ρ = −0.071, p = 0.088, *p-adj* = 0.70).

**Fig. 2.**
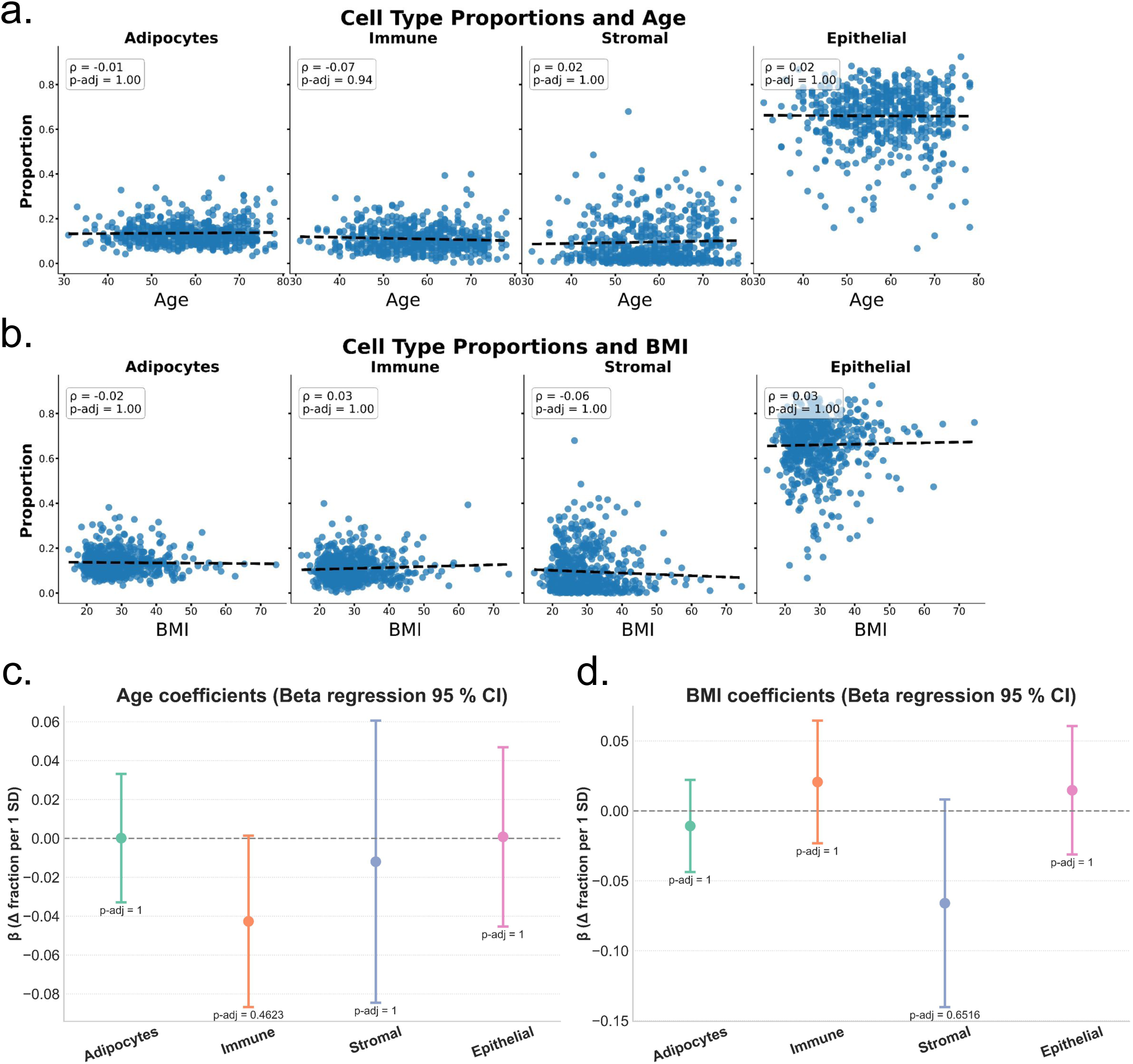
Correlation of cell type proportions in HGSOC tumours by age and BMI. The figure summarizes the relationship of patient age and BMI with proportions of adipocytes, immune cells, stromal cells, and epithelial cells. **a.** Scatterplots with age. Spearman’s ρ and unadjusted p values are displayed in in-plot boxes; fitted least-squares lines illustrate slope direction. Correlation coefficients are small (ρ ∣≤ 0.07∣) and not statistically significant after Bonferroni correction, with largely horizontal trend lines. **b.** Analogous scatterplots for patient BMI, Again, correlation coefficients are small (ρ ∣≤ 0.07∣) and not statistically significant after Bonferroni correction, with largely horizontal trend lines. **c.** Beta-regression coefficients (change in fraction per 1 SD rise in age) with Wald 95% confidence intervals. The dashed horizontal line marks no association. Point estimates for adipocyte, immune and epithelial fractions are near zero; stromal fraction shows a modest negative estimate, but the interval includes zero. All p-values are adjusted for four simultaneous tests, and none meet the significance threshold (annotated beneath each point). **d.** Beta-regression results for BMI mirror the age findings: coefficients are small, confidence intervals span zero, and all multiplicity-adjusted p-values are non-significant. BMI; body mass index.

Beta-regression, which simultaneously adjusts for age and BMI and models residual heteroskedasticity via the precision sub-model, likewise detected no association after adjusting for multiple comparisons (Fig. 2c-d). The strongest association was a decrease in immune-cell proportion with increasing age (β = −0.046 per SD, 95 % CI −0.090 to −0.002, p = 0.039), which was not statistically significant after Bonferroni adjustment (*p-adj* = 0.31). The estimated associations with adipocyte, stromal, and epithelial fractions were small and imprecise (|β| ≤ 0.065; all *p-adj* ≥ 0.68). Collectively, these analyses suggest that age and BMI explain little of the inter-patient variability in tumor adipocyte, immune, stromal, or epithelial content within this cohort.

### Analysis of cell composition by race, residual disease, tissue site, stage, and HGSOC transcriptomic subtype

To examine potential differences in cell fractions by race, we compared the predicted cell proportions’ values of Black and White patients with Welch two-sample *t*-tests, applying Bonferroni correction for the four simultaneous comparisons: one per cell group. We found no significant differences in the cell group fractions between Black and White patients after Bonferroni correction (Fig. 3a). The mean adipocyte content was almost identical (0.136 in Black vs 0.135 in White; *p-adj* = 1.0). The immune (0.111 vs 0.107), stromal (0.090 vs 0.098), and epithelial (0.663 vs 0.661) fractions were also similar (all *p-adj* = 1.0). Thus, in this cohort, estimated differences in adipocyte, immune, stromal, or epithelial abundance do not differ by race.

**Fig. 3.**
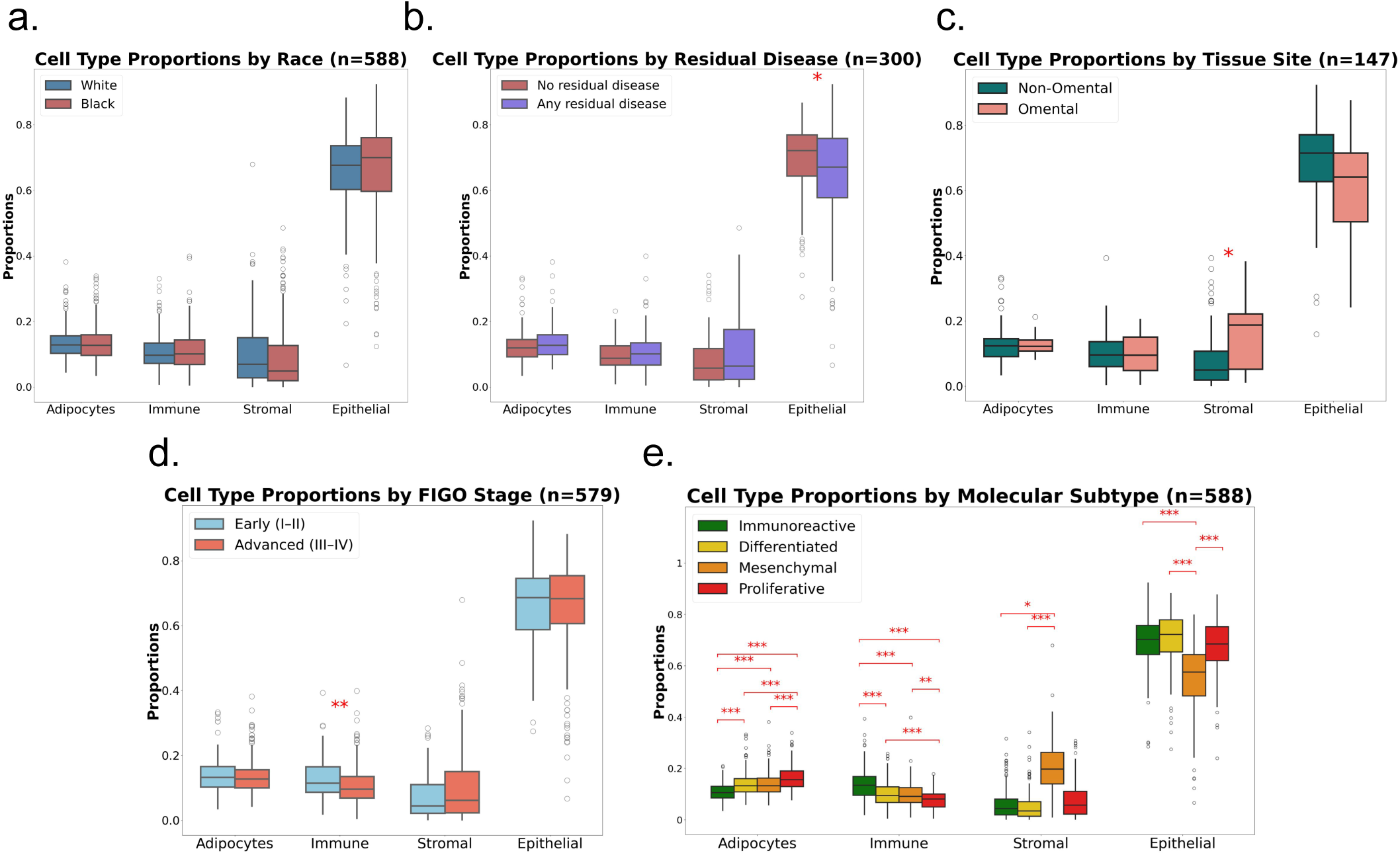
Cell type proportions by race, residual disease, tissue site, stage, and HGSOC transcriptomic subtype Box-and-whisker plots display the distributions of adipocyte, immune, stromal and epithelial cell type proportions (y-axis) for each comparison group (x-axis); horizontal lines mark medians. Red asterisks indicate Bonferroni-adjusted significance levels (**\*** *p-adj* < 0.05; **\*\*** *p-adj* < 0.01; **\*\*\*** *p-adj* < 0.001). **a. Race:** Proportions in White (blue) and Black (red) patients are very similar; no cell type proportion differs (Welch t-test). **b. Residual disease:** Tumours with any macroscopic residual disease (purple) show a modest stromal increase and epithelial decrease compared with completely resected tumours (red), but the difference does not reach the multiple-testing threshold in the stromal case; other compartments are unchanged (Welch t-test). **c. Tissue site:** Tissue samples of omental origin (pink) exhibit significantly higher stromal content than non-omental samples (green, ***), while adipocyte, immune and epithelial fractions remain similar (Mann–Whitney U). **d. FIGO Stage:** Advanced-stage tumours (III–IV, red) have reduced immune infiltration *(**) and elevated stromal proportion* (does not reach significance after Bonferroni adjustment) relative to early-stage disease (I–II, blue). Adipocyte and epithelial fractions do not differ by stage (Welch t-test). **e. Molecular subtype:** ConsensusOV subtypes show distinct cellular compositions in each with distinct signatures, with several highly statistically significant contrasts (one-way ANOVA on logit-transformed fractions followed by Tukey Honestly Significant Difference (HSD) contrasts).

Because residual tumor after surgery could signal a more fibrotic milieu, we tested whether any macroscopic residual disease was associated with cell fractions by contrasting “no residual” versus “any residual” tumours with Welch *t*-tests and Bonferroni adjustment (Fig. 3b). Adipocyte (mean 0.128 vs 0.135; *p-adj* = 1.0) and immune fractions (0.098 vs 0.107; *p-adj* = 0.34) showed no detectable difference. Stromal content was modestly higher in tumors with residual disease (mean 0.107 vs. 0.081; raw p = 0.019), but the difference did not reach Bonferroni significance (*p-adj* = 0.077). The epithelial content showed a significant association with no residual disease presence (mean 0.693 vs. 0.651; raw p = 0.026), with tumours with no residual disease after surgery showing more epithelial content. While epithelial cells are more frequent in tumours with no residual disease and stromal cells may accumulate in tumours with sub-optimal cytoreduction surgery (i.e., residual disease presence), the other two macro compartments were not associated with residual disease status.

We reasoned that the adipose-rich omentum might harbour a distinct microenvironment, which we hypothesized would contain a higher proportion of adipocytes compared to non-omental samples. We compared omental samples with all other anatomic sites using Mann-Whitney tests for each fraction and Bonferroni-corrected for multiple testing. We found only the stromal compartment differed significantly by tissue site (Fig. 3c). Omental samples contained a larger stromal fraction (mean = 0.157) than non-omental tumours (mean = 0.0925; Mann-Whitney *p-adj* = 0.0495). Epithelial content was lower in omental tissue (mean 0.614 vs 0.663) but the difference was not statistically different after Bonferroni correction (*p-adj* = 0.718), and neither adipocyte (0.1295 vs 0.1352, *p-adj* = 1.00) nor immune fractions (0.099 vs 0.109, *p-adj* = 1.00) showed any site-specific difference.

To determine whether disease presentation is associated with differences in the TME, we contrasted early (FIGO I–II) and advanced (III–IV) tumors via Welch *t*-tests on each fraction with Bonferroni correction (Fig. 3d). Advanced tumors contained fewer immune cells (mean 0.105 vs 0.129; Welch *t p-adj* = 0.0072) and more stromal cells (mean 0.096 vs 0.074; *p-adj* = 0.0579), although this did not reach Bonferroni significance, than early-stage tumors. Adipocyte (mean 0.133 vs 0.142; *p-adj* = 0.9073) and epithelial fractions (mean 0.666 vs 0.655; *p-adj* = 1.0) were similar by stage.

We evaluated whether the four TCGA-style consensusOV [6] transcriptomic subtypes, which encode distinct cellular ecosystems, had different distributions of the four macro fractions.

ConsensusOV [6] assignment of 588 tumors categorized them into Immunoreactive (n = 173), Differentiated (n = 180), Mesenchymal (n = 132), and Proliferative (n = 103). We used omnibus one-way ANOVAs on logit-transformed fractions, followed by Tukey HSD (Honestly Ssignificant Difference) contrasts, to test the relationships.

All four macro fractions differed across the four subtypes, each retaining statistical significance after Bonferroni correction: adipocyte (F = 43.193, *p-adj* = 1.28 × 10^⁻24^), immune (F = 33.335, *p-adj* = 2.69 × 10^⁻19^), stromal (F = 5.526, *p-adj* = 3.83 × 10⁻^3^) and epithelial (F = 50.935, *p-adj* < 1.19 × 10⁻^28^). Further Tukey post-hoc contrasts showed a coherent pattern (Fig. 3e):

- **Immunoreactive tumors** were immune-rich and carried the lowest adipocyte load.
- **Mesenchymal tumors** were stromal-enriched, with the lowest proportion of epithelial/tumor cells.
- **Proliferative tumors** were dominated by adipocyte cells and had the sparsest immune infiltrate.
- **Differentiated tumors** had intermediate levels of all three non-epithelial compartments and similar epithelial proportions as immunoreactive and proliferative.

Each subtype displayed a unique signature, affirming that deconvolution-derived composition estimates captured differences between subtypes, further supporting the idea that the transcriptomic subtypes reflect differences in cell composition.

Finally, we examined whether race modifies subtype-specific cell patterns, since it had been previously observed that Black patients had a higher percentage of the Immunoreactive subtype compared to White patients [12]. We tested this by fitting two-way ANOVAs (subtype, race and their interaction) to the same logit-transformed fractions. Adding race as a second factor did not materially change this picture: two-way ANOVA (subtype × race) showed that subtype remained a strong main effect for all four fractions (all *p-adj* ≤ 1.1 × 10⁻²), whereas race alone had no statistically significant influence (all *p-adj* = 1). Interaction terms were non-significant for any compartment (all *p-adj* ≥ 0.75). These results suggest that tumor subtypes are associated with TME composition.

## Discussion

HGSOC is a leading cause of gynecologic cancer mortality, with limited improvement in survival despite advances in treatment [1, 10, 26]. Our study adds a novel dimension by highlighting intratumoral adipocyte content as an independent predictor of poor survival in HGSOC. While past mechanistic studies have shown that adipocytes promote tumor progression through lipid transfer, cytokine signaling (e.g., IL-6, IL-8), and metabolic reprogramming [18, 20, 21, 27], our analysis is among the first to quantify adipocyte abundance directly from bulk RNA-seq and examine associations with patient outcomes. A 10% increase in adipocyte proportion corresponded to an approximate 40% increase in the hazard of death. We also observed an approximate 20% reduction in the hazard of death associated with a 10% increase in immune cells. These findings remained after accounting for age, BMI, stage, race, and residual disease, and mutually adjusting for cell types. The association between increased adipocyte proportion and reduced survival, independent of immune cell proportions, may align with evidence that omental adipocytes promote ovarian cancer metastasis and fuel tumor growth in adipose-rich niches such as the omentum.[19].

We expected that tumors from the omentum would have higher adipocyte content, but in the 11 omental tumors in our study, adipocyte content was not higher than in tumours from other tissue sites. However, omental tumors had a higher proportion of stromal cells (fibroblasts and endothelial cells). While this could be due to the small number of omental tumours included in our analysis, this might be explained by adipocyte plasticity. It is possible that omental adipocytes do not simply accumulate as fat cells in the tumour, but instead actively differentiate into fibroblastic, stromal cells [27–29]. If this is the case, it would contribute to stromal enrichment while diminishing adipocyte proportions in the omentum [18, 29]. Also, since the *Schildkraut* samples were specifically selected for tumor cellularity, it is possible that this reduced the proportion of adipocytes captured in the omental samples. This also suggests that traditional TME analyses with the “missing cell type” problem that excluded adipocytes [30] may have missed the influence of a key cell population.

Previous RNA-seq deconvolution efforts often overlooked adipocytes due to technical limitations in single-cell protocols, which struggle to capture large, lipid-rich cells which behave differently in solution than other cell types. Using the methodological innovation of incorporating snRNA-seq from adipose tissue into a composite deconvolution reference enabled the inclusion of adipocytes. This approach addresses a critical blind spot in standard deconvolution workflows [30] and can serve as a model for future TME profiling in other cancer types.

Our findings position intra-tumoural adipocyte burden as a potentially actionable biomarker. As this signal is extractable from routine bulk RNA-seq data, it could augment risk stratification and help tailor therapies. Moreover, adipocyte-rich tumors may exhibit vulnerabilities to interventions targeting lipid metabolism, adipokine signaling, or adipocyte-fibroblast transition pathways [17].

Beyond these associations, we also examined whether the transcriptional subtypes were related to cell-type proportions. Historically, transcriptomic subtyping using bulk RNA-seq aimed to identify intrinsic tumor categories to inform prognosis and therapy [3, 4, 7, 9, 10], but reproducibility challenges and shifting subtype definitions have led to debate about the biological basis of these classifications. Recent studies have proposed that these subtypes largely reflect TME or cell composition variation captured in bulk through RNA-seq, rather than stable tumor-intrinsic transcriptional programs [12, 13, 15]. This study supports and extends that paradigm.

Through RNA-seq deconvolution of HGSOC tumours, we show that adipocyte proportions vary across transcriptomic subtypes, which have distinct cellular compositions. Immunoreactive tumors were enriched for immune cells, mesenchymal tumors for adipocytes and stroma, and proliferative tumors for epithelial content. These insights mirror conclusions from prior deconvolution and single-cell studies, including recent work by Geistlinger et al. [13] and Hippen et al. [15], and help resolve inconsistencies in previous subtype frameworks. Beyond these subtype-specific cell-type driven patterns, our analysis found that race exerts no main or subtype-specific effect and no detectable subtype-specific influence.

In summary, this study lies at the intersection of several emerging threads in HGSOC research. We introduce adipocytes as a potentially key and quantifiable determinant of prognosis and reinforce the view that bulk transcriptomic subtypes are manifestations of TME composition. We believe this study also demonstrates the value of integrated single-cell and single-nucleus data in resolving cell-type representation gaps in RNA-seq deconvolution [24]. By highlighting the clinical importance of tumor adipocytes, our work opens new directions for both mechanistic investigation and translational application in ovarian cancer.

## Author contributions

Conceptualization: A.I., C.S.G.

Methodology: A.I., C.S.G., S.C.H., J.A.D., L.G.

Software: A.I., G.Y.A.

Formal Analysis: A.I., G.Y.A., L.G.

Data Curation: A.I., G.Y.A., J.M.S., J.A.D., L.C.P., N.R.D., J.R.M.

Writing - Original Draft: A.I.

Writing - Review & Editing: A.I., C.S.G., S.C.H., G.Y.A., J.A.D., L.G., L.C.P., N.R.D., J.R.M.

Visualization: A.I., C.S.G., J.A.D., L.G.

Funding Acquisition: C.S.G., S.C.H., J.M.S.

## Declaration of interests

The authors declare no competing interests.

## Declaration of generative AI and AI-assisted technologies in the writing process

During the preparation of this work, the authors used ChatGPT (5) to improve the grammar and flow of the paragraphs. After using these tools, the authors reviewed and edited the content as needed and take full responsibility for the content of the publication.

## Methods

### Dataset Acquisition and Molecular Subtyping

We leveraged RNA-seq raw counts and the associated metadata from the HGSOC dataset named *Schildkraut* [12]. The dataset contains data from 588 patients total [12], and is composed of two cohorts: one of Black patients and one of White patients. The clinical data used in this study are presented in Table 1.

For molecular subtyping, we log10-transformed the RNA-seq counts [log10(x + 1)], and applied the consensusOV package (version 3.19) [6] using the *get.subtypes* function with default settings and method = “consensus” to assign the four canonical TCGA subtypes: Immunoreactive, Differentiated, Mesenchymal, and Proliferative.

### Single-Cell and Single-Nucleus Reference Data Preprocessing

For the cell reference used for bulk deconvolution, we used 8 HGSOC scRNA-seq samples (GEO accession GSE217517) [31], which do not contain adipocyte cell types. We therefore appended our cell reference with snRNA-seq-derived adipocytes from both subcutaneous and visceral datasets (GEO accession GSE176171) [32]. To incorporate both data modalities in a single reference, we filtered out a list of cross-modality differentially expressed genes, as shown in previous work [24]. This list was previously found to be an effective way of improving deconvolution accuracy and robustness in a previous study by our group, and is publicly available on GitHub [33].

For both the adipocyte snRNA-seq datasets and the HGSOC scRNA-seq datasets, cell types were assigned in the original publication [31, 32]. In HGSOC dataset, cells labeled as *Unknown1* or *Unknown2* were excluded to avoid ambiguous identities. Remaining cell types were then used for the deconvolution references.

For preprocessing quality control, we used the same cells (all others are filtered out) as used in the original publication of the HGSOC data [31]. For the adipocyte snRNA-seq data, in addition to the original publication’s quality control filtering [32], we used a stringent per nucleus quality control to make sure only the highest quality adipocytes are included in the final reference. We excluded nuclei exhibiting > 150 total mitochondrial reads, > 10 % mitochondrial read fraction, < 200 detected genes (debris), or > 4,500 detected genes (accounting for doublets/multiplets). All preprocessing steps are reproducible and available in the GitHub repository [34].

### Deconvolution of HGSOC RNA-seq Bulks

For deconvolution, we used the BayesPrism framework implemented in InstaPrism for computational efficiency [35]. BayesPrism is a probabilistic framework that leverages the expression profiles of the cell reference to deconvolve bulk mixtures that has been shown to outperform other frameworks [36]. We executed the InstaPrism pipeline for 5,000 iterations for each bulk. The bulk and single-cell data (cell reference) are input as raw counts, as expected by the deconvolution method. The deconvolution output consisted of estimated cell type proportions for each bulk sample.

### Cox Proportional Hazard Models for Survival Analysis

The *Schildkraut* cohort included only Black and White individuals. Because there were only three Hispanic patients (0.5% of all patients) and two patients with unknown ethnicity, we only considered race (and not ethnicity) as a variable in our analyses. Of the 588 individuals, we excluded 13 patients with unknown BMI and nine patients with unknown FIGO stage, for a total of 566 patients included in the main survival analyses. Because 49% of patients were missing data on residual disease, we also performed analyses restricted to patients with known residual disease status and adjuvant chemotherapy treatment information.

For the restricted model, after excluding patients with unknown BMI and FIGO stage, 566 patients remained. Restricting that cohort to those with known residual disease status yielded 293 patients. We then excluded the 7 patients who were known not to have received adjuvant chemotherapy, leaving 286 patients. Of those patients, we further restricted the analysis to patients whose adjuvant treatment information was available, resulting in a final restricted-model cohort of 262 patients.

Left-truncated survival models were fit, with time at risk defined from study enrollment (interview date) to all-cause mortality or censoring, rather than from time of diagnosis, as done previously to account for immortal time bias [37]. All CPH models assessed associations with cell type proportion as the exposure and mortality as the outcome, and were stratified by FIGO Stage (i.e., strata = Stage, early (FIGO I/II) or advanced (III/ IV)). The CPH model assumptions were checked for each model using the same package used to fit the models, Lifelines 0.30.0 [38], and all models passed, unless noted. The main models were adjusted for age, BMI, and race.The time axis for both models was days from interview to last follow-up or death.

### Kaplan Meier Curves for Missing Treatment Information

The *Schildkraut* data includes a subset of patients with no treatment information who either had no surgery or for whom surgery status was not available (Table 1). To evaluate the potential survival impact of missing treatment information, Kaplan-Meier survival analysis was conducted using the same Lifelines package [38]. Patients were categorized into two groups: those with available data for all two treatment-related variables (adjuvant therapy and residual disease status) and those missing all. Kaplan-Meier curves were plotted for these two groups, separately by stage: advanced stage (FIGO I /II) and early-stage (FIGO III/IV). Furthermore, we repeated the analysis as detailed above separated by race Black patients and White patients.

Survival distributions were compared visually, with survival probability plotted as a function of time in days. This analysis assessed whether the absence of treatment metadata might introduce systematic bias in downstream survival modeling, and curves are shown in Supplemental Fig. 5. There was no indication of survival differences between patients with missing information compared to patients with present information, in both all patients and separated by race.

### Continuous covariates: Age and BMI Analysis

Continuous covariate analyses were performed on the 575 *Schildkraut* tumors that had age and BMI data (13 patients with unknown BMI were not included). The four macro-fractions (adipocytes, stromal, immune, and epithelial) were derived as described above. Each fraction was correlated with both age and BMI using Spearman’s ρ, giving eight tests in total. Two-sided p-values were adjusted with a Bonferroni correction (α = 0.05/8 ≈ 0.0063).

#### Beta-regression

Age and BMI were standardized to z-scores so that regression coefficients represent the change in fraction produced by a one-SD increase in the covariate (subtract the average, divide by the population standard deviation). Because β-regression can requires numbers that are between 0 and 1, values were clipped to the open interval (ε, 1 - ε) with ε = 5 × 10^-^⁴ to satisfy β-regression assumptions.

For each macro fraction, we fitted a logit link β-regression (*statsmodels* version 0.14.0) [39], which first converts the fraction into logit space so the linear part of the model can take any value. The model included two predictors, the z-score standardized age (Age-z) and BMI (BMI-z), in both the mean (μ) and precision (ϕ) sub-models, plus an intercept. Without an intercept, the model would force the mean (or precision) to zero when both predictors are zero, and is non-applicable for proportions. Wald 95% confidence intervals, raw p-values, and standardised coefficients (β) were extracted. The resulting eight p-values (four fractions x two predictors) were adjusted using a Bonferroni correction. Model diagnostics (Pearson residuals vs fitted, studentised QQ-plot, leverage/Cook’s D) were inspected for all fractions. Diagnostic plots showed symmetric, homoscedastic residuals with only a handful of moderate-influence points (Cook’s D ≤ 0.33); exclusion of those points did not materially change any coefficient. These plots can be seen in the GitHub repository notebook *analysis_proportions_vs_bmi_and_age.ipynb* [34].

### Clinical covariate and cell group proportion analysis

For the *Schildkraut* HGSOC cohort we merged the cleaned clinical data with cell-fraction proportion estimates obtained from bulk-RNA-seq deconvolution (see above), yielding 588 tumors with both clinical metadata and cell fractions.

Fine-grained deconvolution cell-type proportion outputs were collapsed into the four high-level macro fractions of adipocytes, stromal, immune, and epithelial by summing the relevant single-cell clusters (see Supplemental Fig. 6 for mapping).

#### Race

For each macro fraction we compared distributions between Black and White patients using Welch’s unequal-variance *t*-test. Direct proportions are used in this test because only two groups were compared and Welch’s test handles heteroscedasticity directly.

#### Residual disease

For the 300 samples whose residual disease status was recorded, surgical outcome codes were collapsed to two classes: No residual tumor (0 mm) and any residual tumor (< 1 cm, ≥ 1 cm, or size-unknown). For each macro fraction, we ran a Welch two-sample *t*-test comparing the two residual-disease classes (unequal variance).

#### Tissue site

Tumors were classified as Omental (primary or metastatic samples labelled “Omentum”, n=16) or Non-Omental (all ovary, fallopian-tube, mixed adnexal and “Other” sites, n=131). Using the same collapsed macro fractions (adipocytes, stromal, immune, epithelial), we compared the two site classes with a two-sided Mann-Whitney U test, chosen because several macro fractions showed non-normal distributions (Fig. 4c).

#### Stage

Tumors with recorded stage (n = 579) were dichotomised into early disease (FIGO I-II) and advanced disease (FIGO III-IV). Group means were compared with Welch’s unequal-variance *t*-test on the raw proportion scale for the four macro fractions (Fig. 3d).

For all comparisons, four simultaneous tests (one per macro fraction) were Bonferroni-adjusted, with *p-adj* < 0.05 considered significant. Box-plots display group distributions, with red asterisks marking Bonferroni-significant contrasts.

### Transcriptomic Subtype Analysis

We investigated cell-type proportion differences first across transcriptomic subtypes alone and then using models that included race and a subtype-by-race interaction. After merging consensus subtype calls with the bulk-deconvolution table, 588 tumors had complete data (Immunoreactive = 175, Differentiated = 179, Mesenchymal = 130, Proliferative = 104; Black = 272, White = 316). Because comparisons involved four groups and, in the second analysis, an interaction term, we used analysis-of-variance (ANOVA) models, which assume roughly normal, homoscedastic residuals. The raw proportions (bounded 0-1) were right-skewed and showed unequal variances among subtypes, so each macro fraction (adipocytes, immune, stromal, epithelial) was variance-stabilized with the logit transformation: logit (p) = log[p / (1 - p)], where p is the cell group proportion. This extra step was unnecessary in earlier two-group comparisons, which were handled with Welch t-tests or Mann–Whitney U tests that are robust to heteroscedasticity without transformation.

For the one-way analysis, we ran four omnibus ANOVAs, one per logit-transformed fraction. The resulting p-values were Bonferroni-corrected; fractions with *p-adj* < 0.05 were followed by Tukey HSD pair-wise contrasts to pinpoint specific subtype differences.

For the two-way analysis, we fitted ordinary least-squares models of the form logit (p) ∼ Subtype + Race + Subtype:Race. Type-II sums-of-squares tables provided F statistics and p-values for the Subtype main effect, Race main effect, and their interaction, accommodating the unequal cell counts across subtype × race combinations. P-values from the two-way ANOVA (four fractions × three effects) were adjusted using a Bonferroni correction. Box-plots were then annotated with significance stars for fractions or pair-wise contrasts meeting the α = 0.05 threshold (Fig. 3e).

## Data Availability

The clinical data and the unnormalized RNA-seq count data for the *Schildkraut* datasets are available upon request. All additional data that support the findings of this study (deconvolution reference) are freely available online for download. The HGSOC data used can be downloaded through Gene Expression Omnibus GSE217517 [31], and all snRNA-seq adipocyte data can be downloaded through GSE176171 [32]. All data details and download links can be easily accessed in the GitHub repository [34].

## Code Availability

The code developed for this study is available at our GitHub repository [34] under BSD 3-Clause License, and a Zenodo archive will be created once this article is accepted. The repository includes all necessary scripts and a README file with setup instructions and usage guidelines. For updates or assistance, users can refer to the repository or open an issue for queries. Our aim is to support transparency and reproducibility in computational research through this open-access resource.

## Acknowledgements

This work was supported by NCI of NIH (R01 CA200854 to J.A. Doherty and J.M. Schildkraut, R01 CA142081 to J.M. Schildkraut, R01 CA076016 to J.M. Schildkraut, R01 CA188943 to J.M. Schildkraut, and R01 CA237170 to C.S. Greene and J.A. Doherty). Research reported in this publication utilized the High-Throughput Genomics and Cancer Bioinformatics Shared Resource at Huntsman Cancer Institute at The University of Utah and was supported by NCI of the NIH under award number P30CA042014. The data used in this manuscript were generated from participants recruited via the Rapid Case Ascertainment (RCA) method and the North Carolina Central Cancer Registry (CCR). It is a research study data collection method used by the University of North Carolina Lineberger Comprehensive Cancer Center supported by P30CA016086 and the University Cancer Research Fund of North Carolina. RCA is a collaboration between UNC Lineberger, the North Carolina Central Cancer Registry (CCR), and participating hospitals in North Carolina.

## Notes

https://github.com/greenelab/deconvolution_sc_sn_comparison

### Competing Interest Statement

The authors have declared no competing interest.

