## Supplementary Materials for "Deconvolved tumor adipocyte proportions and high grade serous ovarian carcinoma survival"

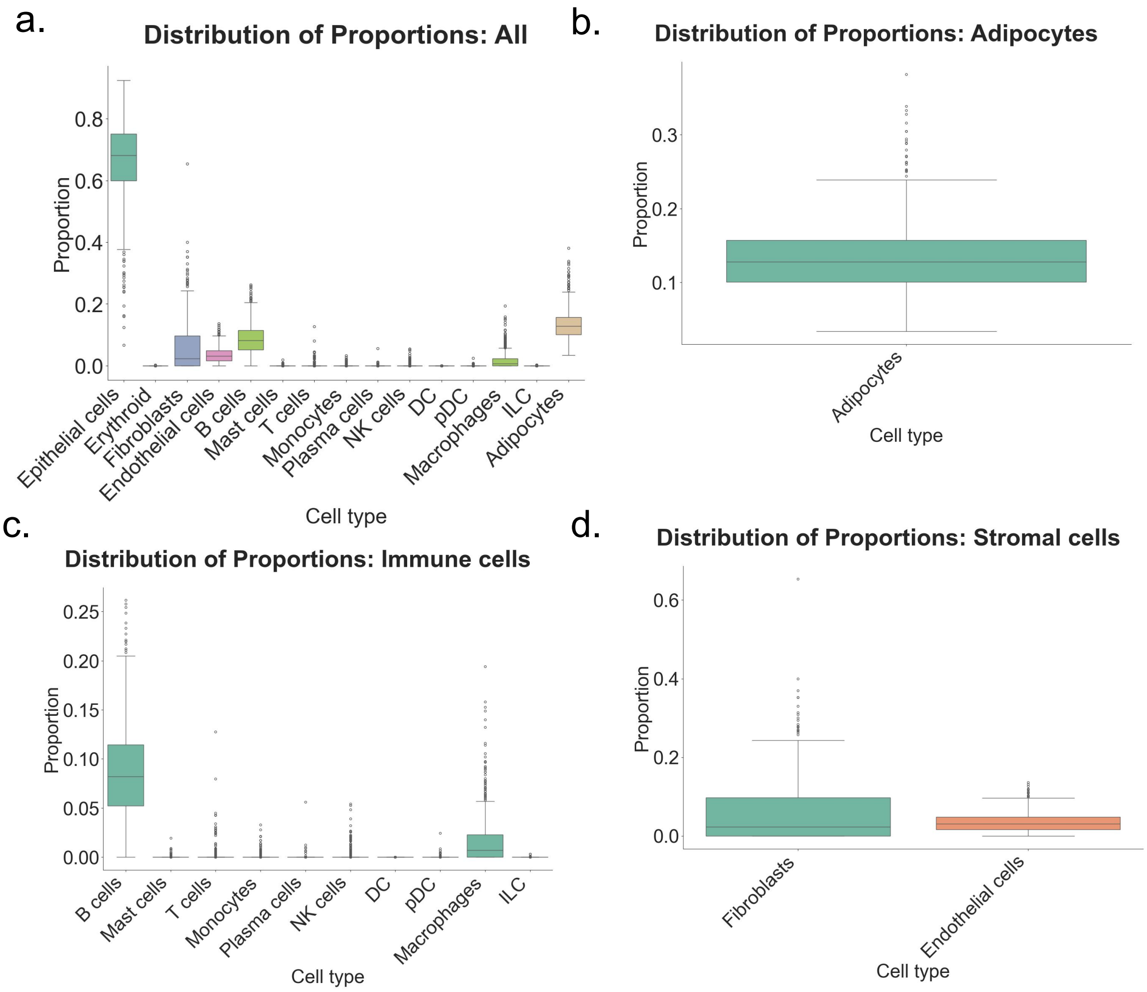


**Supplemental Fig. 1**

**Distribution of the predicted cell type proportions by bulk RNA-seq deconvolution**

The figure shows boxplots of the distributions of the predicted proportions across all 588 *Schildkraut* samples. a. The distributions of all cell types b. the distributions of adipocytes, c. the distribution of the immune cells (summed to form the immune macro-fraction), and the distribution of the stromal cells (summed to form the macro-fraction).

NK cells; Natural killer cells. DC; Dendritic cells. PDC; Plasmacytoid dendritic cells. ILC; Innate lymphoid cells.

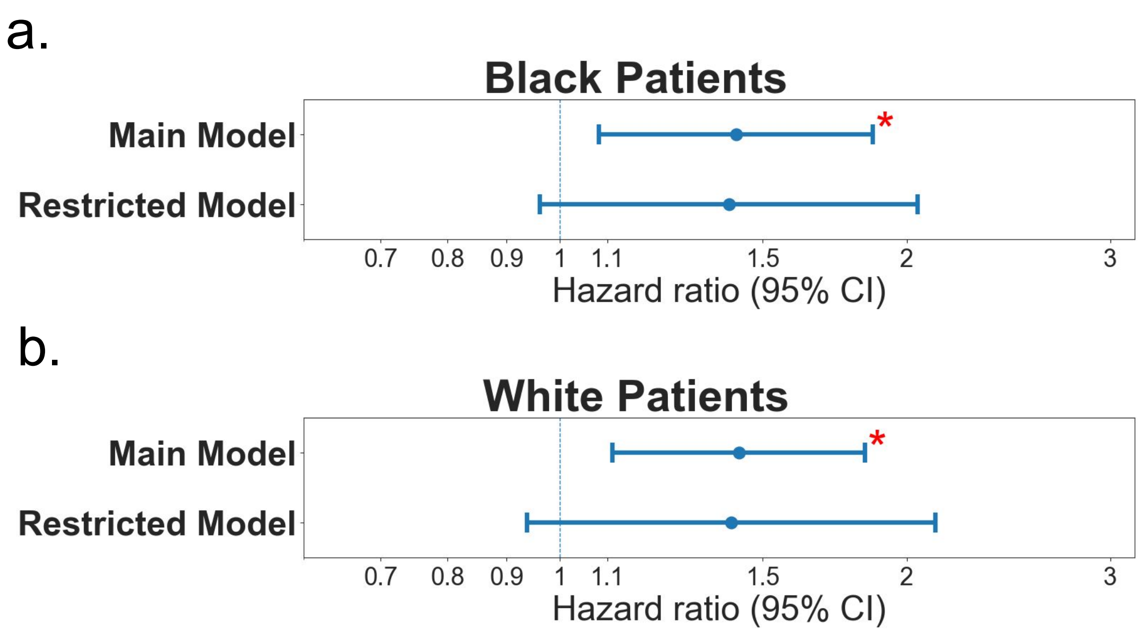


**Supplemental Fig. 2**

**Adipocyte proportion and high-grade serous ovarian cancer survival separately by race**

The figure illustrates the forest plots of the survival analysis parallel to Fig. 1, but with results stratified for Black and White patients. Panel a shows the main (n = 263) and the restricted model (n = 151) for Black patients only, and panel b. shows the main (n = 303) and the restricted model (n = 111) for White patients. All forest plots show the HR from the CPH models on the x-axes. Red asterisks indicate statistically significant associations (* $p < 0.05$; ** $p < 0.005$; *** $p < 0.0005$). The HR = 1 vertical line at HR = 1 serves as a reference for no effect. CPH; Cox Proportional hazard. HR; Hazard Ratios.

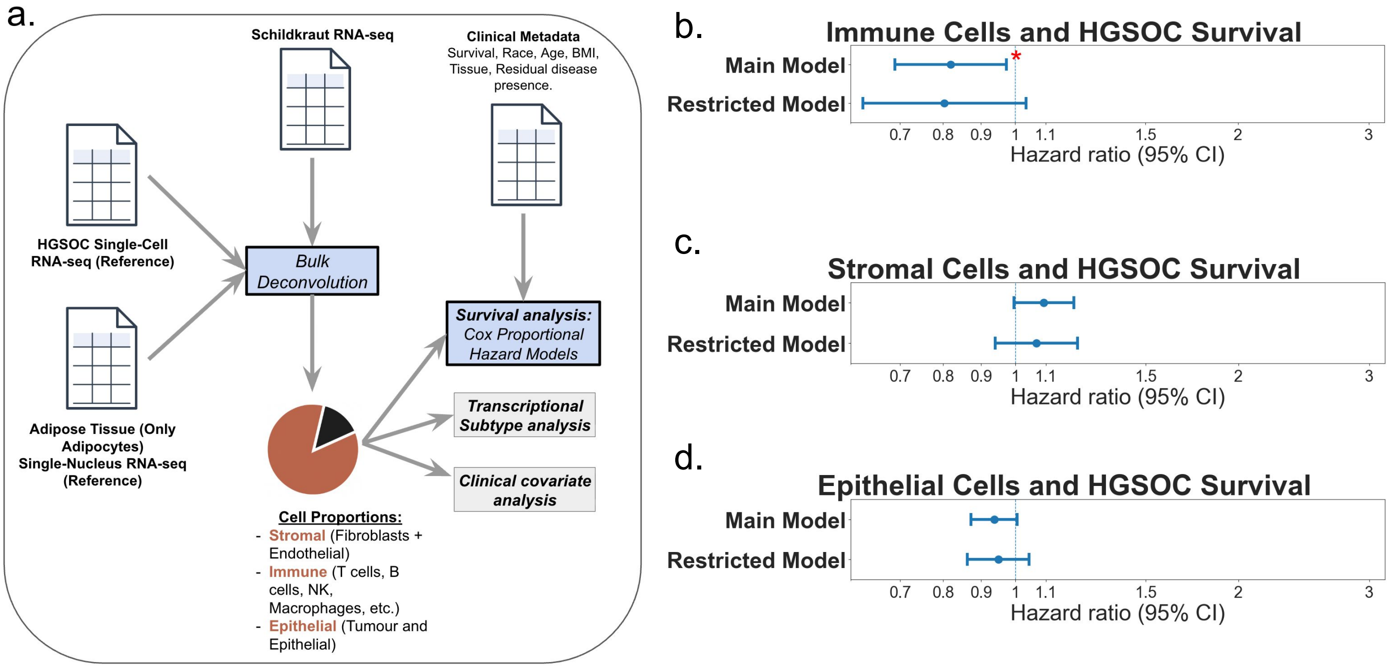


**Supplemental Fig. 3**

**Immune, stromal, and epithelial proportion quantification and association with high-grade serous ovarian cancer survival**

The figure parallels Fig. 1 but focuses on subsequent analysis for other TME compartments. **a.** Fine-grained cell types predicted by bulk deconvolution (see Fig. 1a) were collapsed into three macro groups: immune (T, B, NK, myeloid, etc.), stromal (fibroblasts + endothelial), and epithelial. For each tumor, the component fractions were summed and multiplied by ten so that a one-unit change equals a 10 percentage-point rise. These scaled fractions, together with clinical covariates, were entered into stage-stratified, CPH models. **b-d.** Forest plots of HRs from the CPH analyses. Each subpanel shows two models: the **main** model (top) controls for age, BMI, and race; the **restricted model** (bottom) additionally controls for residual disease presence, with n = 566 and n = 262 patients, respectively. Red asterisks indicate statistically significant associations (* $p < 0.05$; ** $p < 0.005$; *** $p < 0.0005$). The dashed vertical line marks HR = 1 (no association).

CPH; Cox Proportional hazard. HR; Hazard Ratios. BMI; body mass index.

a.

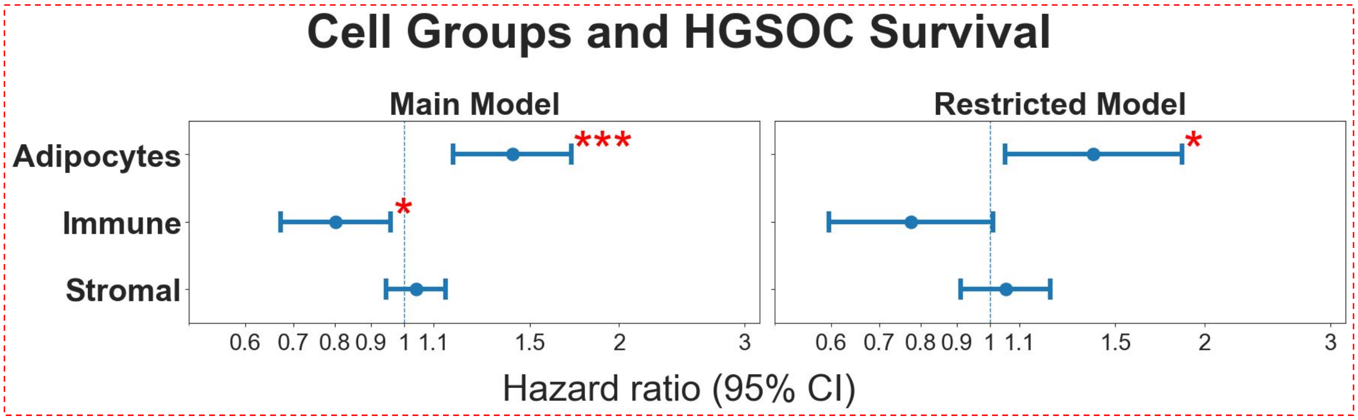


*Note: Both the main and restricted CPH models showed a violation of the proportional hazards assumption for the Adipocytes variable (Schoenfeld test p = 0.0289 and 0.0543 respectively). These models are marked with dashed red lines. Due to this violation, these model's estimates are considered unstable and are presented for transparency only; combined Black and White patients Cox model results are used for primary inference.*

**Supplemental Fig. 4**

**Sensitivity analysis of adipocyte, immune, and stromal (removed epithelial) proportions and association with high-grade serous ovarian cancer survival**

The figure focuses on a sensitivity analysis of cell proportion relationship with survival while taking all cell populations into account in one model. Fine-grained cell types predicted by bulk deconvolution (see Fig. 1a) were collapsed into three macro groups: immune (T, B, NK, myeloid, etc.), stromal (fibroblasts + endothelial), and epithelial. For each tumor, the component fractions were summed and multiplied by ten so that a one-unit change equals a 10 percentage-point rise. These scaled fractions, together with clinical covariates, were entered into stage-stratified, CPH models. We removed the epithelial compartment to serve as the reference cell proportion for these models. **a.** Forest plots of HR from the CPH analyses in the main model and the restricted model (n = 566 and n = 262). Red asterisks indicate statistically significant associations (* $p < 0.05$; ** $p < 0.005$; *** $p < 0.0005$). The dashed vertical line marks HR = 1 (no association). Note that both models do not pass the CPH assumptions test (Schoenfeld) and are only shown for transparency rather than inference.

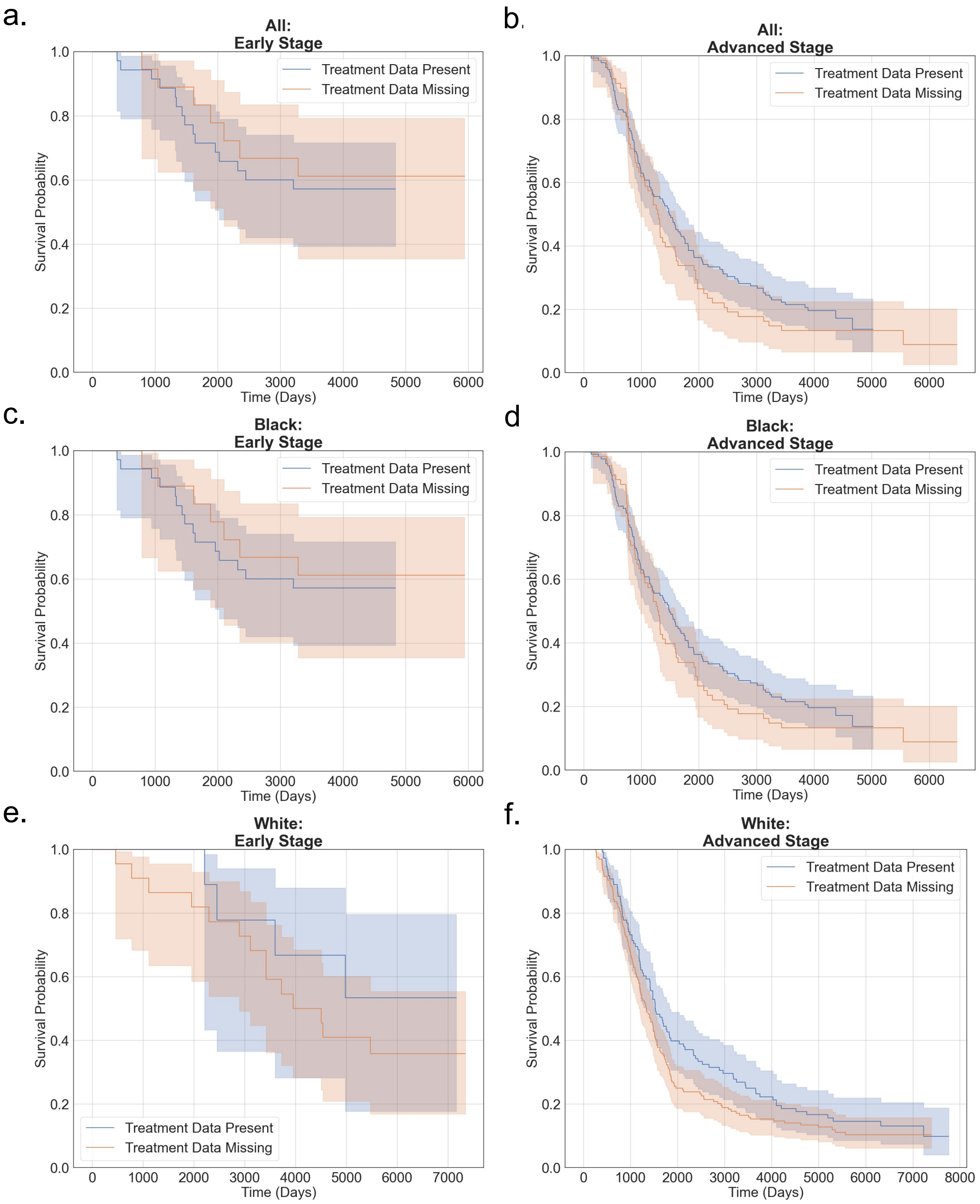


**Supplemental Fig. 5**

**Kaplan-Meier survival by main and restricted models.**

The left column shows early-stage (FIGO I-II) cases, and the right column shows advanced-stage (FIGO III-IV) cases. Panels a and b contain data on all patients included in the main analyses, panels c and d contain data for all Black patients, and panels e and f contain data for all White patients. Orange curves represent the 566 patients included in the main models (with some missing data on receipt of chemotherapy and residual disease), and blue curves represent the 262 patients included in restricted models because they have data on residual disease. Shaded bands denote 95% confidence intervals. In both stage strata the survival trajectories overlap extensively, and log-rank tests are non-significant, indicating that absence of treatment data is not associated with differential overall survival.

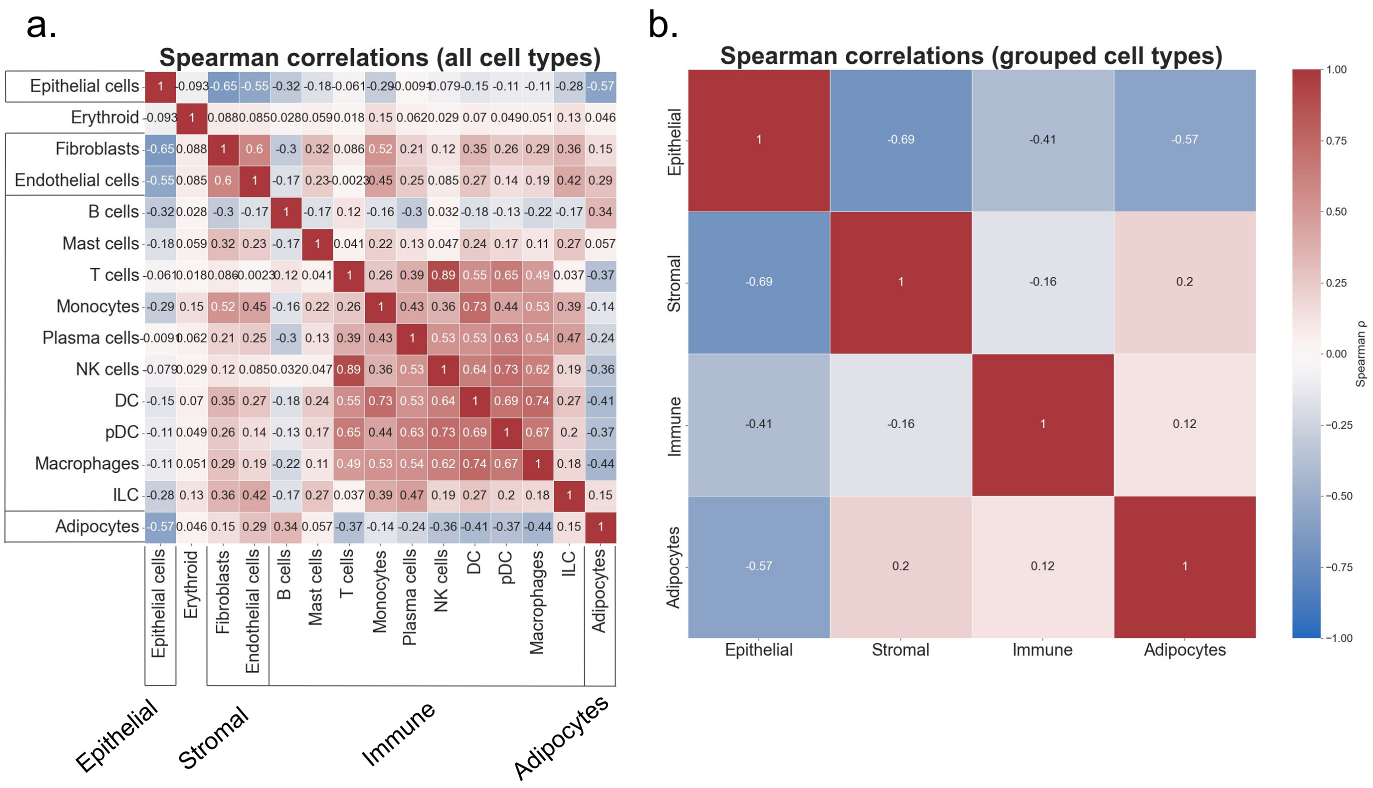


**Supplementary Fig. 6**

**Spearman rank correlations among deconvolved cell fractions**

Heat-map of pair-wise Spearman ρ values for all individual cell types (n = 588 tumors). Blocks along the axes indicate the defined four macro compartments used later: epithelial, stromal, immune and adipocyte. Within-compartment correlations are generally positive (deep red), whereas cross-compartment correlations are mostly negative or weakly positive. Color scale ranges from –1 (dark blue) to +1 (dark red), with numeric ρ values superimposed. a. Shows the assigned cell types and their mapping to the subsequent groups in b.

NK cells; Natural killer cells. DC; Dendritic cells. PDC; Plasmacytoid dendritic cells. ILC; Innate lymphoid cells.
